# High-content phenotypic screen to identify small molecule enhancers of Parkin-dependent ubiquitination and mitophagy

**DOI:** 10.1101/2022.09.30.509127

**Authors:** Roberta Tufi, Emily H. Clark, Tamaki Hoshikawa, Christiana Tsagkaraki, Jack Stanley, Kunitoshi Takeda, James M. Staddon, Thomas Briston

## Abstract

Mitochondrial dysfunction and aberrant mitochondrial homeostasis are key aspects of Parkinson’s disease (PD) pathophysiology. Mutations in PINK1 and Parkin proteins lead to autosomal recessive PD, suggesting that defective mitochondrial clearance via mitophagy is key in PD etiology. Accelerating the identification and/or removal of dysfunctional mitochondria could therefore provide a disease-modifying approach to treatment. To that end, we performed a high-content phenotypic screen (HCS) of ~125,000 small molecules to identify compounds that positively modulate mitochondrial accumulation of the PINK1-Parkin-dependent mitophagy initiation marker p-Ser65-Ub in Parkin haploinsufficiency (Parkin ^+/R275W^) human fibroblasts. Following confirmatory counter-screening and orthogonal assays, we selected compounds of interest that enhance mitophagy-related biochemical and functional endpoints in patient-derived fibroblasts. Identification of inhibitors of the ubiquitin-specific peptidase and negative regulator of mitophagy USP30 within our hits further validated our approach. The compounds identified provide a novel starting point for further investigation and optimisation.

## INTRODUCTION

Mitochondrial dysfunction is a prominent pathological feature of both sporadic and familial Parkinson’s disease (PD) and has long been implicated in disease etiology ^1–8^. Monogenic PD implicates mitochondria as central to disease pathogenesis ^9^. Mutations in mitochondrial PTEN (phosphatase and tensin homologue)-induced kinase 1 (PINK1; encoded by *PARK6*) and Parkin (encoded by *PARK2/PRKN*), cause autosomal recessive early onset PD ^2,5,8,10,11^. PINK1 and Parkin act in concert to identify and subsequently remove damaged mitochondria by selective autophagy (mitophagy), being, respectively, sensor and amplifier proteins of the mitophagy process ^3,4,12^. Importantly, mitochondrial dysfunction and reduced rates of mitophagy are evident in sporadic PD ^13–20^, and variants in genes associated with mitochondrial function are associated with PD risk ^21^, again linking mitochondrial health and clearance processes to PD pathophysiology. Taken together, enhancing mitochondrial clearance by mitophagy is a promising disease-modifying strategy in PD.

The PINK1-Parkin mitochondrial quality control system is a well-studied pathway in PD. In healthy cells, the kinase PINK1 is targeted to mitochondria and N-terminally translocated to the inner mitochondrial membrane (IMM). Sequential proteolysis and proteasomal degradation maintain low basal levels of PINK1 protein ^22^. Mitochondrial damage, typically presenting as loss of mitochondrial membrane potential (MMP), stabilises the active PINK1 protein on the outer mitochondrial membrane (OMM) ^23^. Stabilised PINK1 auto-phosphorylates, homodimerises, and then phosphorylates serine 65 (Ser65) residues of ubiquitin present at the mitochondrial surface under basal conditions ^24–27^. The E3-ubiquitin ligase Parkin binds phospho-Ser65-ubiquitin (p-Ser65-Ub) and is phosphorylated by PINK1 on the homologous Ser65 residue of its own ubiquitin-like (UBL) domain ^26,28,29^. These binding events and post-translational modifications release the auto-inhibitory conformation of Parkin, stabilising an active conformation ^29,30^. Parkin further ubiquitinates mitochondrial surface proteins ^31^, facilitating a feed-forward amplification loop of substrate ubiquitination, phosphorylation, and Parkin recruitment to tag damaged mitochondria ^12,29,32,33^. Mitochondrial surface proteins Miro and mitofusin1/2 are targeted for proteasomal degradation to modulate mitochondrial dynamics ^31,34,35^, and poly-ubiquitinated OMM proteins act as receptors for autophagic adaptors and permit autophagosome formation for removal by the lysosome ^36^.

As PINK1 is the only recognised Ser65-ubiquitin kinase ^28,37^, mitochondrial stress-induced phosphorylation of ubiquitin (p-Ser65-Ub) represents a key biomarker of mitophagy initiation. Here, we established a target-agnostic, high-content phenotypic screen (HCS) to identify small molecule enhancers of p-Ser65-Ub in a familial PD human fibroblast carrying a heterozygous *PRKN* R275W point mutation. We screened ~125,000 compounds from the Eisai compound library and, following hit confirmation and counter-screening, identified compounds displaying desirable activity and drug-like properties to commence a drug discovery campaign. These compounds were further prioritised based on pharmacological profile and assessment of compound-driven functional *en masse* mitochondrial clearance. Biochemical screening of hits led to identification of putative inhibitors of the mitophagy negative regulator USP30 (a ubiquitin specific peptidase/deubiquitinase; DUB), thus validating our approach. We believe that the shared mechanistic basis between HCS assay endpoint and disease phenotype creates robust predictive power for therapeutic development, enabling identification of novel small molecule modifiers of mitophagy in PD.

## RESULTS

### Development of a p-Ser65-Ub high content phenotypic assay

To confirm our methodology and the contribution of PINK1-Parkin signalling to the accumulation of p-Ser65-Ub, we first quantified mitochondrial stress-induced phosphorylation of ubiquitin in control, Parkin ^−/−^ and PINK1 ^−/−^ SH-SY5Y cells via immunocytochemistry (ICC; Fig. 1A). A marked increase in p-Ser65-Ub levels – indicative of PINK1-Parkin pathway activation – was observed in isogenic control cells following incubation with FCCP or antimycin A/oligomycin A (AA/O), which was significantly reduced in Parkin ^−/−^ cells, and abolished in PINK1 ^−/−^ cells (Fig. 1A), suggesting p-Ser65-Ub is a reliable and specific reporter of PINK1-Parkin function and is both detectable and quantifiable using ICC.

**Fig. 1.**
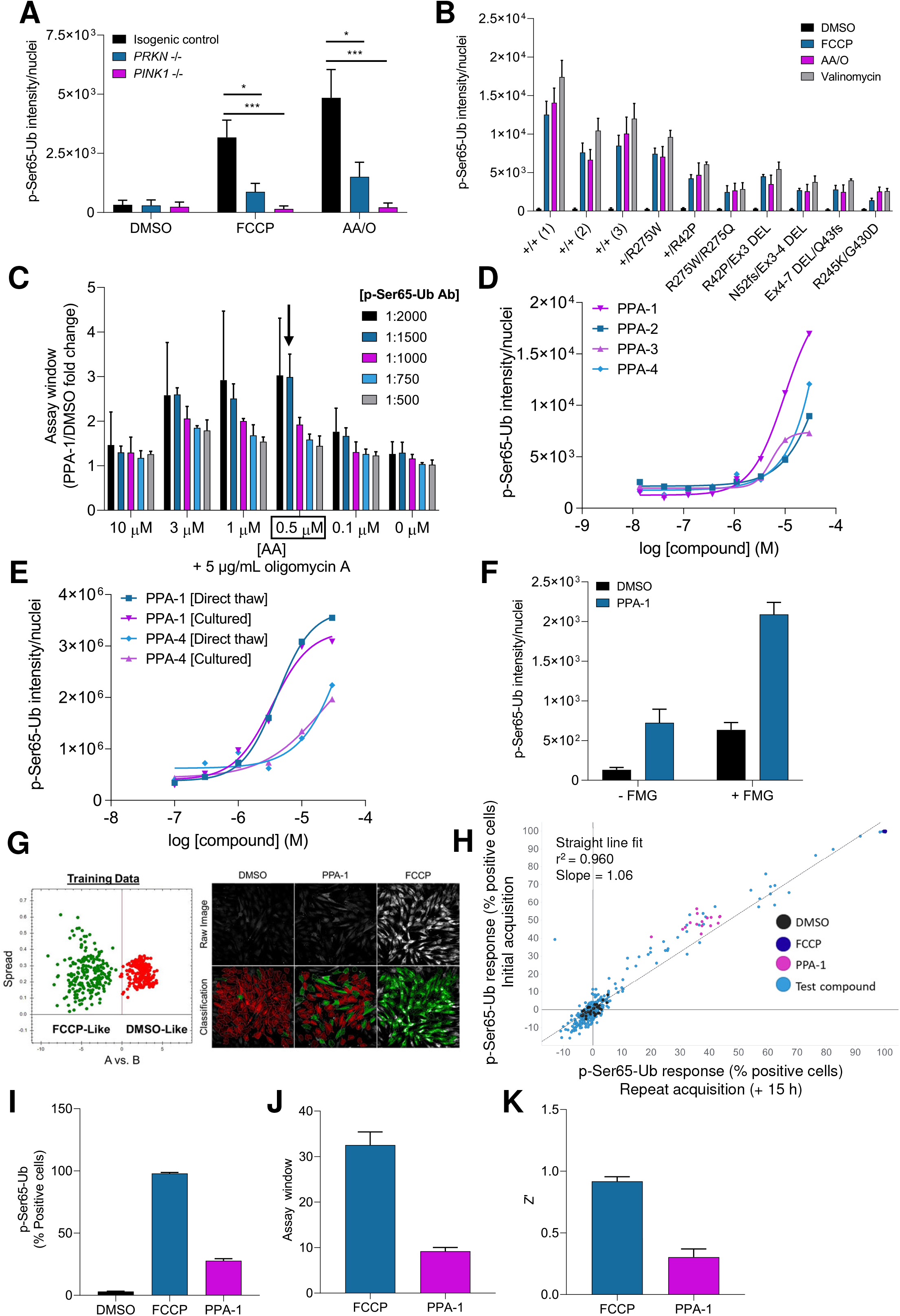
Development of a p-Ser65-Ub high-content phenotypic assay. **A**, p-Ser65-Ub response following 2 h treatment with DMSO (vehicle), 10 μM FCCP or 10 μM antimycin A/5 μg/mL oligomycin A (AA/O) in *PINK1^−/−^* and *PRKN^−/−^* SH-SY5Y cells. **B**, p-Ser65-Ub response following 2 h treatment with DMSO (vehicle), 10 μM FCCP, 5 μM AA/O or 200 nM valinomycin in patient-derived healthy control fibroblasts [+/+ (1), (2) and (3)] and those carrying *PRKN* mutations (Supplementary Table 1). **C**, p-Ser65-Ub response following 2 h treatment with increasing concentration of AA/O (0.1 to 10 μM) in Parkin ^+/R275W^ fibroblasts. p-Ser65 ICC was performed with increasing concentration of anti-p-Ser65-Ub antibody, with 20 μM PPA-1 (2 h pre-incubation) used to determine maximal assay window. Black box and black arrow indicate the stimulus and antibody concentration, respectively, selected for the HCS. **D**, p-Ser65-Ub concentration-response curves of PPA-1-4 (2 hr pre-incubation) under optimised HCS assay conditions (0.5 μM AA/O) in Parkin ^+/R275W^ fibroblasts. **E**, p-Ser65-Ub concentration-response curves of PPA-1 and −4 (2 h pre-incubation), in both cultured and directly thawed Parkin ^+/R275W^ fibroblasts under HCS assay conditions. **F**, p-Ser65-Ub response in Parkin ^+/R275W^ fibroblasts in the presence and absence of the Fluoromount G (FMG) anti-fade reagent, following treatment with 20 μM PPA-1 (2 h pre-incubation) in HCS assay conditions. **G**, *Left panel*, phenotype A versus phenotype B illustration scatterplot describing spread of image texture properties of the training data set obtained using PhenoLOGIC. Each coloured dot reflects a single cell following treatment with DMSO (red) or FCCP (20 μM; green) under HCS assay conditions in Parkin ^+/R275W^ fibroblasts. 6 fields and over 200 individual cells were selected per condition. Trained algorithm was used for subsequent HCS image analysis. *Right panel*, representative images and application of trained algorithm of p-Ser65-Ub response in Parkin ^+/R275W^ fibroblasts following treatment with DMSO, 20 μM PPA-1 and 20 μM FCCP. Cells classified as ‘DMSO-Like’ (red) and ‘FCCP-Like’ (green) phenotypes. **H**, Scatterplot, correlation coefficient (r^2^) and line of best fit for p-Ser65-Ub normalised response between initial and repeat (15 h post-processing) image acquisition of the same plate. **I**, p-Ser65-Ub response of positive (PPA-1 and FCCP; both 20 μM) and negative controls (DMSO) in Parkin ^+/R275W^ fibroblasts under HCS assay and analysis conditions. **J**, Determination of the assay window using data derived in **I**. **K**, Determination of the Z’ using data derived in **I**. For **A**, error bars represent s.d. (*n* = 3 independent experiments). \**P* < 0.05 and \*\*\**P* < 0.001; two-way ANOVA with Bonferroni’s multiple comparisons compared to isogenic control cells. For **B**, error bars represent s.d. (*n* = 3 independent experiments). For **F**, data are mean ± s.d. (*n* = 1, 8 technical replicates per condition). For **I** to **K**, data are mean ± s.d. (*n* = 1, 6 technical replicates per condition).

Previously, we identified patient-derived familial PD fibroblasts which demonstrate mitophagy deficits and decreased p-Ser65-Ub accumulation, with a clear genotype-phenotype relationship determined by allelic number of *PRKN* mutations ^38^. Furthering these observations, we profiled p-Ser65-Ub responses in an expanded panel of patient-derived Parkin mutant fibroblasts (Fig. 1B). We found significant deficits in p-Ser65-Ub accumulation associated with severity of *PRKN* genotype across a range of mitophagy-inducing stimuli, with homozygous or compound heterozygous mutants being most affected (Fig. 1B). These data suggest a strong relationship between *PRKN* status and stress-induced p-Ser65-Ub accumulation and, together with evidence of mitochondrial dysfunction in sporadic PD ^13–20^, provide rationale for the development of a p-Ser65-Ub endpoint assay to identify molecules augmenting PINK1-Parkin signalling to normalise mitochondrial deficits.

We next developed a high-content phenotypic screening assay to classify compounds capable of enhancing p-Ser65-Ub accumulation. Parkin ^+/R275W^ fibroblasts have a reduced p-Ser65-Ub response compared to common variant controls (Fig. 1B) and, importantly, we have previously shown enhanced p-Ser65-Ub accumulation in these cells following USP30 inhibition, suggesting that p-Ser65-Ub can be modulated via pharmacological intervention of the PINK1-Parkin pathway ^38^. As such, a high-content imaging assay for p-Ser65-Ub was established in Parkin ^+/R275W^ fibroblasts, with the aim of analysing small molecule-mediated normalisation of mitophagy deficits in a familial PD cellular background. In addition to p-Ser65-Ub, HSP60 and LAMP1 were included as markers of mitochondrial and lysosomal compartments, respectively.

Following automation and miniaturisation of the p-Ser65-Ub ICC assay to a 384-well plate format, we optimised assay conditions. We selected AA/O treatment over 2 hour incubation as the acute mitophagy-inducing stimulus for our assay. We titrated AA and antibody concentration to determine the largest assay window in the presence of putative Parkin activator-1 (PPA-1; derived from patent WO 2018/023029) (Fig. 1C). A three-fold change in p-Ser65-Ub with PPA-1 was measured in 0.5 μM AA (in the presence of constant 5 μg/mL oligomycin A) using the fluorophore conjugated p-Ser65-Ub (E2J6T)-Alexa647 at 1:1500 dilution, resulting in a large assay window (Fig. 1C). Higher antibody concentrations failed to show a p-Ser65-Ub change, and lower concentrations (1:2000 dilution) produced greater signal variability (Fig. 1C). At lower AA/O concentrations a small p-Ser65-Ub response was observed, and high AA/O concentrations revealed a saturated p-Ser65-Ub response that could not be enhanced by PPA-1, each limiting the assay window (Fig. 1C). To confirm our assay was fit-for-purpose, putative Parkin activators (PPA-1-4) were tested under optimised assay conditions, obtaining a concentration-response for all four compounds (Fig. 1D). PPA-1, which produced the largest p-Ser65-Ub response, was selected as a mechanistically relevant positive control for the HCS.

To improve efficiency and consistency throughout the screen, we identified that direct thaw of bulk cryopreserved cells produced equivalent pharmacological profiles for PPA-1 and PPA-4 compared to cells maintained in culture (Fig. 1E). As such, cells were batch prepared to generate over 1×10^9^ cells, enough to complete primary and counter-screening, allowing us to ensure cells were passage-matched throughout the HCS and to minimise impact of variation in culture conditions. The addition of the anti-fade reagent Fluoromount G (FMG; diluted 1:1 in D-PBS) following antibody incubation resulted in improved p-Ser65-Ub signal intensity and preservation (Fig. 1F) and was used routinely.

An image analysis method was developed using the Harmony PhenoLOGIC machine learning plug-in (PerkinElmer) to score cells as either p-Ser65-Ub positive or p-Ser65-Ub negative, based on the image texture phenotype of the positive (FCCP) and negative (DMSO) controls (Fig. 1G). FCCP was selected as the positive control given the reliable enhancement of p-Ser65-Ub in these cells (Fig. 1B). In the presence of FMG and using the PhenoLOGIC analysis, assay readout was stable across 15 hours at room temperature (the longest duration plates were waiting to be imaged in a single run; Fig. 1H).

Finally, we validated FCCP and PPA-1, our two positive controls in the assay (Fig. 1I–K). FCCP produced a large p-Ser65-Ub signal (Fig. 1I) and assay window (Fig. 1J), and a Z-Prime (Z’; a statistical measure of assay robustness ^39^) between 0.8-0.9, indicating a high-quality assay (Fig. 1K). PPA-1 produced a lower p-Ser65-Ub signal, assay window, and Z’ (Fig. 1I-K) but was more mechanistically relevant (a putative mitophagy enhancer rather than a mitochondrial toxin). On this basis, we selected two assay quality control (QC) measures: primary QC was the plate Z’, using the DMSO negative control and FCCP as the primary positive control; secondary QC was the normalised p-Ser65-Ub response to PPA-1, used to confirm run-to-run robustness of assay sensitivity. The p-Ser65-Ub response was normalised based on the negative (DMSO; 0%) and positive (FCCP; 100%) response (percentage positive cells; Fig. 1G). Plates with a Z’ above 0.8 and a secondary QC above 20% normalised response satisfied our screening criteria and were used for further analysis. Based on the response of PPA-1 and the distribution of negative control responses, we selected a ‘hit’ threshold of 15% normalised p-Ser65-Ub response. Thorough optimisation of the p-Ser65-Ub ICC assay protocol allowed us to establish a robust, quality-controlled, and reproducible HCS assay.

### Establishment of a TMRM counter-screen to exclude compounds that affect mitochondrial membrane potential

Mitochondrial function is critically linked to mitochondrial polarisation state. Mitochondrial stress or damage typically presents *in vitro* as loss of MMP triggering PINK1-Parkin mitophagy and the accumulation of p-Ser65-Ub ^12,40^. To understand the relationship between MMP depolarisation and p-Ser65-Ub accumulation in our HCS assay, we examined MMP by tetramethyl-rhodamine methyl ester (TMRM) fluorescence in a well-validated and established imaging assay ^38^. We demonstrated an inverse correlation between TMRM intensity and p-Ser65-Ub accumulation upon treatment with AA/O (Fig. 2A-B). Following 2 hour incubation with AA/O (analogous to HCS conditions), a p-Ser65-Ub response was only observed at concentrations of AA/O which caused a >75% reduction in TMRM fluorescence (Fig. 2B). Our HCS-stimulus condition (0.5μM AA/5 μg/mL oligomycin A) reduced TMRM fluorescence by approximately 60% and did not induce a substantial p-Ser65-Ub percentage positive cell response (Fig 2A-B).

**Fig. 2.**
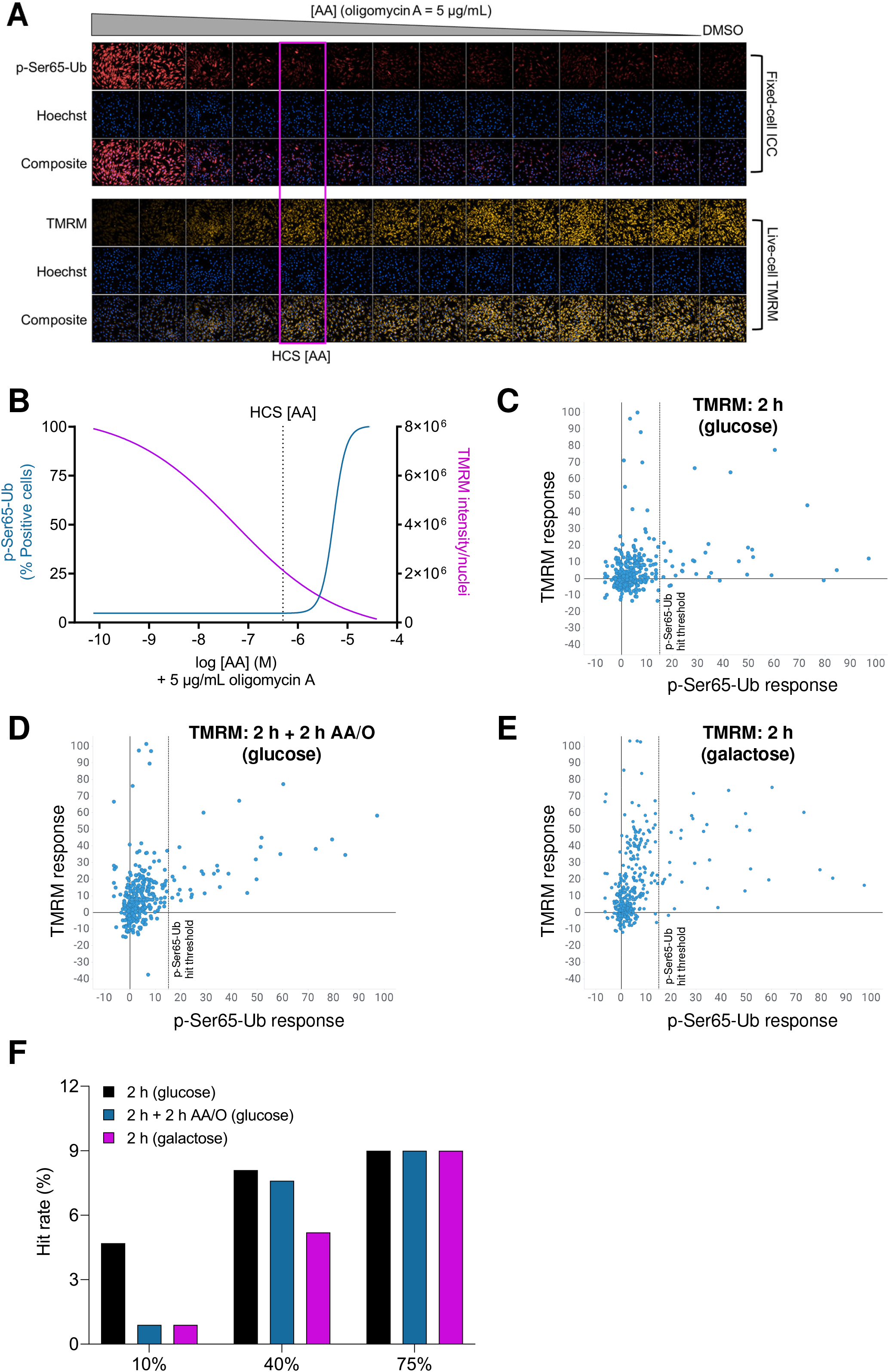
Establishment of a TMRM counter-screen to exclude compounds that affect mitochondrial membrane potential. **A**, Representative images of p-Ser65-Ub and TMRM following 2 h antimycin A (with constant oligomycin A; 5 μg/mL) concentration-response in Parkin ^+/R275W^ fibroblasts. Pink box indicates the antimycin A concentration ([AA]) selected for HCS. **B**, Quantification of TMRM fluorescence intensity and percentage of p-Ser65-Ub positive cells as in **A**. Vertical dotted line indicates the antimycin A concentration ([AA]) selected for HCS. **C-E**, Scatter plots of p-Ser65-Ub normalised response of test compound (blue dots) *versus* TMRM normalised response following: **C**, 2 h treatment with compound in glucose-containing DMEM; **D**, Standard assay conditions of 2 h pre-treatment with compounds followed by 2 h treatment with 0.5 μM AA/O; and **E**, 2 hr treatment with compound in galactose-containing DMEM. In **C**, **D**, and **E**, number of tested compounds (dots) is 344. Black dotted lines mark the 15% p-Ser65-Ub normalised response hit threshold. **F**, Hit rate (percentage of the 344 test compounds which pass both the p-Ser65-Ub threshold (> 15%) and the indicated TMRM threshold) is reported applying different TMRM cut-offs (10, 40 and 75% TMRM normalised response).

Compounds which inherently depolarise MMP or have a cumulative depolarising effect when combined with AA/O would increase p-Ser65-Ub and were designated as ‘hits’ in our HCS assay, despite acting via a nonspecific mechanism. To identify and exclude these undesirable compounds, we developed a TMRM protocol for use as a counter-screening assay (Fig. 2C-F). A representative set of compounds were screened using the p-Ser65-Ub HCS assay, and the TMRM assay was performed in parallel after either 2 hour treatment with compounds (Fig. 2C); 2 hour treatment with compounds followed by 2 hour treatment with AA/O (Fig. 2D), matching the p-Ser65-Ub HCS conditions; or 2 hour treatment with compounds in galactose-substituted media (Fig. 2E); following overnight incubation with galactose, to metabolically force the cells into oxidative phosphorylation instead of glycolysis and make them more sensitive to mitochondrial insult ^41^. Of the 344 compounds tested, 32 compounds hit in the p-Ser65-Ub HCS assay (hit rate = ~10%). When tested in glucose-containing medium without AA/O, most hit compounds had limited effect on TMRM fluorescence (Fig. 2C). However, in the presence of AA/O or in galactose-substituted media, more compounds reduced TMRM fluorescence, indicating an effect on MMP (Fig. 2D-E).

As shown in Figure 2B, a reduction in TMRM of approximately 75% is necessary for a p-Ser65-Ub response in our HCS fibroblast model. However, this may not be a sufficiently harsh threshold for compound selection as MMP depolarisation is a major confounding factor for mitophagy activation and must be avoided. To select a suitable threshold for the TMRM counter-screening assay, we compared hit rates in the three TMRM assay conditions using different exclusion thresholds (Fig. 2F). A relaxed TMRM threshold (40% or 75%) only slightly reduced the 10% hit rate obtained from the p-Ser65-Ub assay alone (Fig. 2F). Using a 10% TMRM cut-off, the hit rate was 5% (Fig. 2F) after 2 h incubation in glucose-containing medium. In contrast, using the same threshold in the TMRM assay in both galactose medium and under HCS conditions (with AA/O) we obtained a 1% final hit rate (Fig. 2F). We therefore selected this stringent 10% TMRM threshold and glucose medium containing AA/O as the finalised assay conditions, as this aligned the TMRM counter-screen with the primary p-Ser65-Ub HCS, permitted precise correlation between endpoints, and produced an estimated final hit rate of 1%, an acceptable outcome from a HCS ^42^.

### Known mitophagy modulators were active in the optimised HCS assay

To validate our screening system, we used our optimised HCS (Fig. 1) and TMRM (Fig. 2) assays to explore the effects of published compounds proposed to affect aspects of mitochondrial and/or lysosomal biology. Forty-five compounds were tested (Fig. 3A), from broad mechanistic classes including proton ionophores, kinase modulators, and proposed mitophagy activators such as PPAs and USP30 inhibitors ^9^. Compounds were tested at low (1:1666 dilution), mid (1:500), and high (1:166) concentration, as determined by the starting concentration of each compound (Supplementary Table 3), and p-Ser65-Ub and TMRM endpoints measured and compared (Fig. 3B-D).

**Fig. 3.**
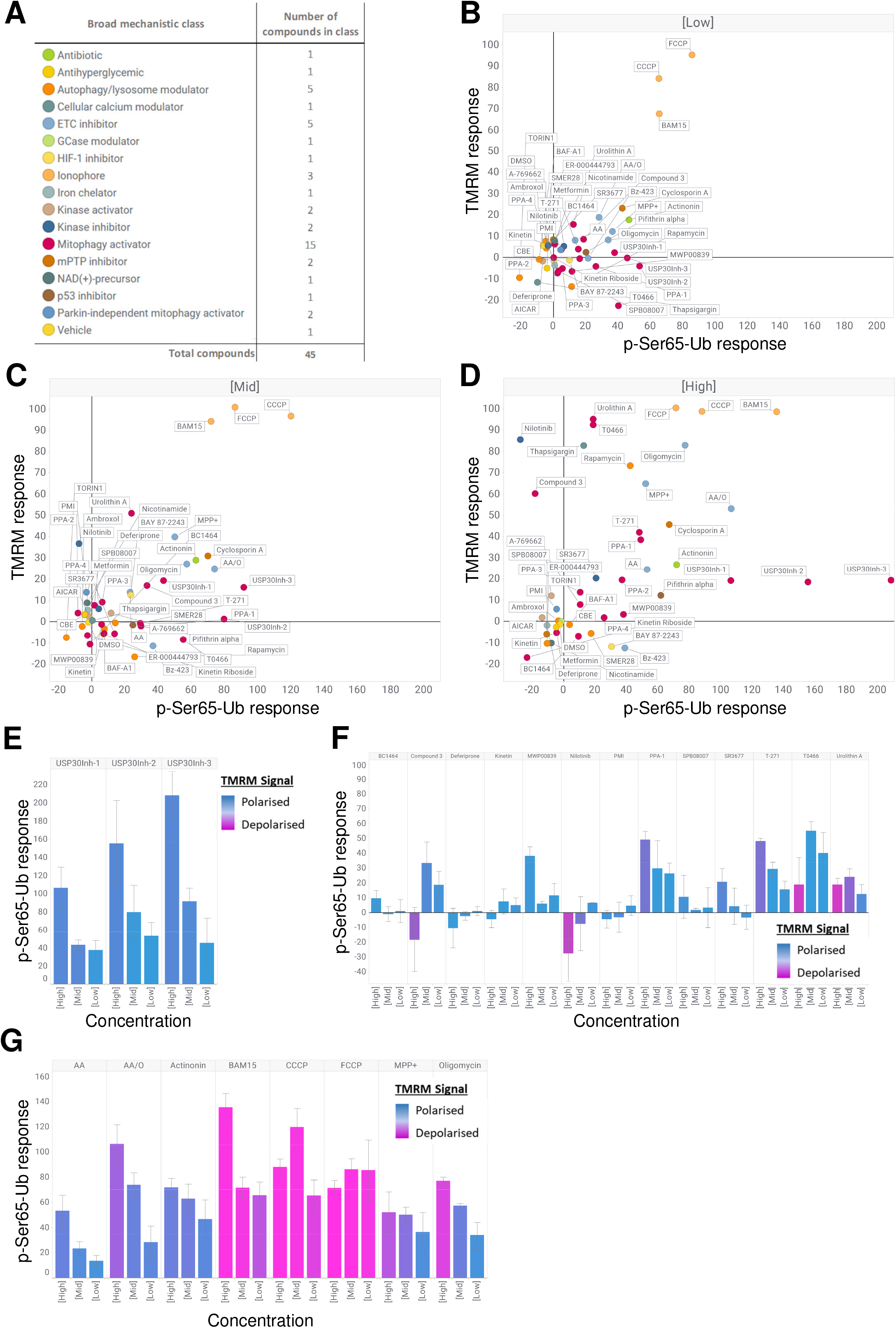
Known mitophagy modulators were active in the optimised HCS assay. **A,** Broad mechanistic class (mechanism of action) of the forty-four literature identified compounds (plus vehicle, DMSO) employed in the proof-of-concept study. **B-D**, Scatter plots of p-Ser65-Ub and TMRM normalised response using HCS primary and TMRM secondary screening protocols for compounds reported in **A** tested at low (**B**), mid (**C**) and high (**D**) concentration. **E**, p-Ser65-Ub and TMRM normalised responses following treatment with low to high concentrations of USP30 inhibitors (USP30Inh-1, −2 and −3), derived from **B-D**. Height of bars indicates p-Ser65-Ub normalised response and colour gradient indicates mitochondrial polarisation state, as determined by TMRM intensity (polarised [blue] – depolarised [pink]). **F**, p-Ser65-Ub and TMRM normalised responses following treatment with low to high concentrations of mitophagy enhancers, derived from **B-D**. Height of bars indicates p-Ser65-Ub normalised response and colour gradient indicates mitochondrial polarisation state, as determined by TMRM intensity (polarised [blue] – depolarised [pink]). **G**, p-Ser65-Ub and TMRM normalised responses following treatment with low to high concentrations of mitochondrial toxins, derived from **B-D**. Height of bars indicates p-Ser65-Ub normalised response and colour gradient indicates mitochondrial polarisation state, as determined by TMRM intensity (polarised [blue] – depolarised [pink]). For **B-G**, data are mean ± s.d. (*n* = 3 independent experiments).

At all concentrations, the protonophores FCCP, CCCP, and BAM15 (6-60 μM) induced strong p-Ser65-Ub responses and a high degree of MMP depolarisation, whereas limited effects on either MMP or p-Ser65-Ub were observed for other compounds at low concentration (Fig. 3B). At increasing (mid and high) concentrations, several compounds demonstrated concentration-dependent effects on MMP and/or p-Ser65-Ub (Fig. 3C and D). The PPAs behaved as expected based on previous observations (Fig. 1), with PPA-1 inducing the strongest p-Ser65-Ub response, although this compound modestly impacted MMP at high concentration (60 μM; Fig. 3D). USP30 inhibitors (USP30Inh-1, 2 and 3) strongly induced p-Ser65-Ub with limited effect on MMP, as observed previously under FCCP-mediated mitochondrial stress ^38^ (Fig 3E).

Several compounds identified in recent studies as enhancers of PINK1-Parkin mitophagy also demonstrate accumulation of p-Ser65-Ub in our assay system, including T-271 (originally identified via a Mito-SRAI screen ^43^), the ROCK2 inhibitor SR3677 (identified from a eGFP-Parkin translocation screen ^44^) and the Fbxo7-FP domain inhibitor, BC1464 (identified from a virtual screen, as a modulator of Fbxo7-PINK1 interactions ^45^) (Fig. 3F). BC1464 only has a modest effect in our assay, however the treatment period employed in our assay system is much shorter than that used in the original paper (2 hour *versus* 16 hour compound incubation) (Fig. 3F). Furthermore, T0466, which induced Parkin translocation and enhanced Parkin-dependent degradation of luciferase-tagged MFN-1 ^46^, increased p-Ser65-Ub at 6 μM and 20 μM, while inducing mitochondrial depolarisation at high concentration (Fig. 3F). We observed no p-Ser65-Ub response using kinetin, a proposed PINK1 activator, however previous experiments with this compound required 24 hour incubation and a technology for cell delivery ^47^ (Fig.3F). MWP00839 at high concentration (60 μM), but not SPB08007, compounds identified using Mito-Timer as enhancers of basal mitophagy ^48^, promoted accumulation of p-Ser65-Ub, with limited effect on MMP (Fig. 3F).

Importantly, our p-Ser65-Ub HCS assay did not identify mitophagy enhancing compounds with mechanism of action independent of the PINK1-Parkin pathway it reports on, including p62-mediated mitophagy inducer (PMI), which enhances mitophagy via p62, downstream of the PINK1-Parkin axis ^49^, and the iron chelator deferiprone ^50^ (Fig. 3F). The c-ABL inhibitor nilotinib ^51^ failed to induce p-Ser65-Ub and demonstrated MMP depolarisation at higher concentration (Fig. 3F). Interestingly, Miro1 reducer Compound 3 ^52^ promoted p-Ser65-Ub accumulation at low-mid concentration in our system (Fig. 3F) but demonstrated a loss of MMP at high concentration. As predicted, mitochondrial toxins increased p-Ser65-Ub, with a heavy dependence on MMP depolarisation (Fig. 3G). Together, these data established our HCS and TMRM counter-screen as fit-for-purpose and demonstrated that we could confidently use these assays to identify specific modulators of PINK1-Parkin biology.

### HCS and counter-screening identified MMP-independent positive modulators of p-Ser65-Ub

To identify novel positive modulators of p-Ser65-Ub within the Eisai compound library we used the finalised HCS conditions (Fig. 4A) to screen approximately 125,000 compounds at a single concentration (~5 μg/mL, with molar concentrations ranging from 20 to 30 μM; Fig. 4B). Cells were pre-treated for 2 hours with either test compound, negative (DMSO), or positive (FCCP or PPA-1) control, then AA/O (final assay concentration [FAC] of 0.5 μM AA and 5 μg/mL oligomycin A) was added as a mild mitophagy-inducing stimulus. Based on the 15% p-Ser65-Ub threshold and assay QC criteria previously established (Fig. 1), we identified approximately 15,000 hits from our primary screen, a hit rate of 12.3% (Fig. 4B-C).

**Fig. 4.**
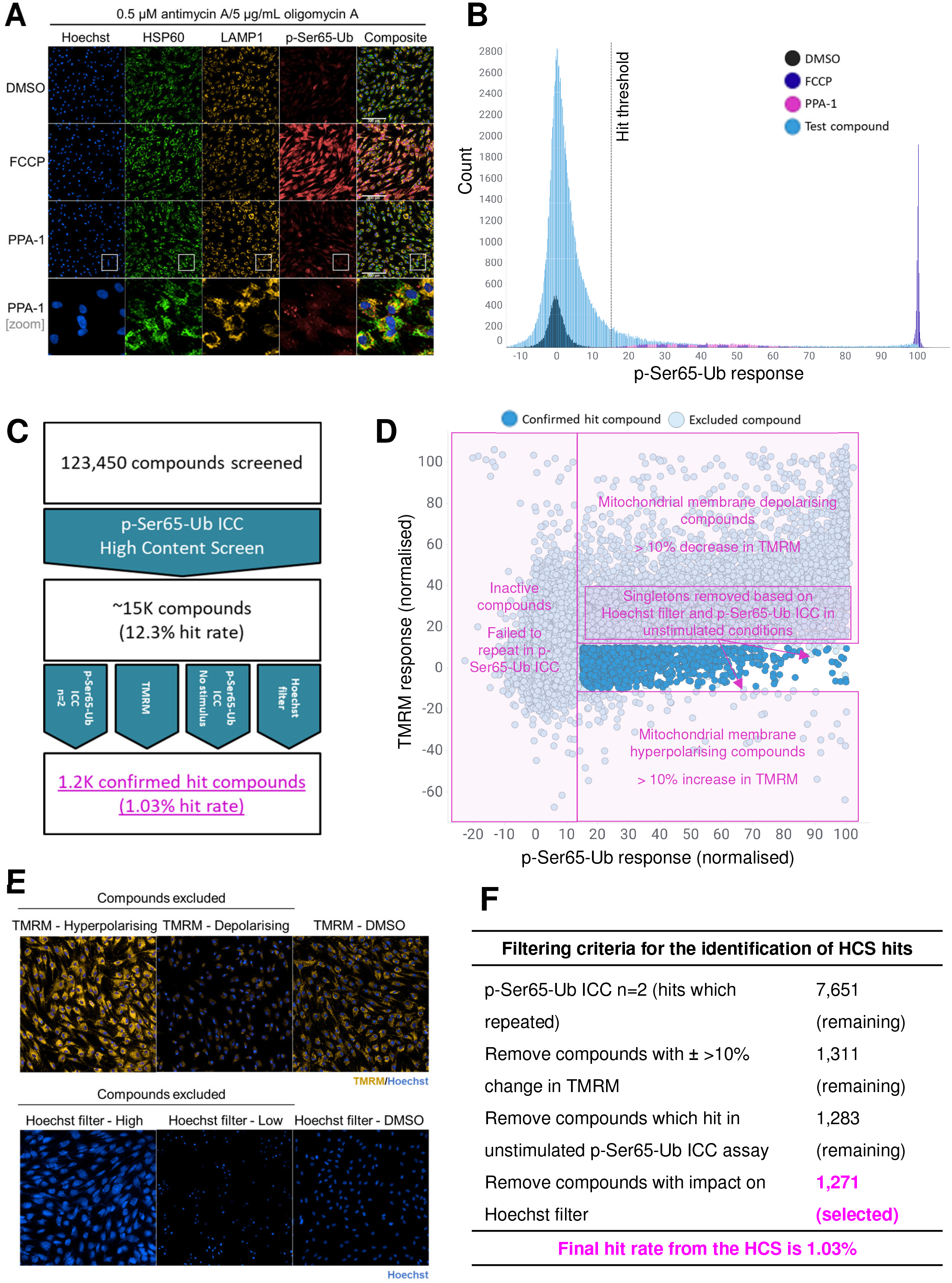
HCS and counter-screening identified positive modulators of p-Ser65-Ub. **A**, Representative images of Parkin ^+/R275W^ fibroblast cells treated with either vehicle (DMSO), 20 μM FCCP, or 20 μM PPA-1 under HCS assay conditions. p-Ser65-Ub (red), HSP60 (green), LAMP1 (yellow) and Hoechst nuclear stain (blue). Scale bar 200 μm. Bottom panels are higher magnification images of the areas in the white boxes. **B**, Primary HCS. Normalised p-Ser65-Ub response after treatment with ~125,000 compounds, each run in singlicate at approximately 20-30 μM, using HCS protocol. Distribution of negative control (DMSO, dark blue), positive control (20 μM FCCP, purple), secondary positive control (20 μM PPA-1, pink), and test compounds (light blue). All data presented passed QC, based on selected criteria (Z’ > 0.8 and secondary QC > 20% response). Dashed line indicates the selected hit threshold of > 15% p-Ser65-Ub response; test compounds with p-Ser65-Ub response above this threshold were selected as hits. **C**, Screening and counter-screening workflow. **D**, Hit compound counter-screening. Scatter plot of p-Ser65-Ub and TMRM normalised response using HCS primary and TMRM secondary screening protocols for ~15,000 compounds identified in primary HCS (20 μM; singlicate; 2 hr pre-incubation). Compounds were excluded based on mitochondrial membrane depolarisation (± 10% change in TMRM response), failure to repeat (p-Ser65-Ub response < 15%), response in unstimulated conditions (p-Ser65-Ub response > 15% in absence of a mitochondrial stressor), and overt effects on Hoechst fluorescence. **E**, Representative images for the counter-screening assays. **F**, Selected filtering criteria for counter-screening assays and number of compounds that passed, derived from data in **D**.

Primary hits were then subjected to counter-screening assays measuring four parameters (Fig. 4C-F). First, we reconfirmed p-Ser65-Ub responses using compound stocks standardised to 20 μM (p-Ser65-Ub n=2). Approximately half (7,651) of the ~15,000 primary hit compounds were confirmed (Fig. 4D and F). We then used our optimised TMRM assay (Fig. 2), excluding both depolarising and hyperpolarising test compounds (Fig. 4D and E). As normalised TMRM response partially correlates with normalised p-Ser65-Ub response (r^2^= 0.416; derived from data in Fig. 4D), exclusion of compounds with substantial depolarising effect resulted in disproportionate removal of compounds with high p-Ser65-Ub response, thus leaving approximately 1,300 hit compounds (Fig. 4D-F). Our third counter-screening parameter was a p-Ser65-Ub assay without a mitophagy-inducing stimulus (without AA/O). Approximately 40 additional compounds which passed the first two filters (Fig. 4F) were eliminated. It worth noting that while these compounds were not deemed useful according to our current screening strategy, they may serve as tool compounds or be potentially useful for other approaches due to their ability to enhance p-Ser65-Ub in the absence of mitochondrial stress. Finally, we applied a filter based on Hoechst fluorescence, excluding compounds which increased or decreased Hoechst fluorescence intensity by greater than two standard deviations from the Hoechst mean intensity of all compounds (Fig. 4E-F). The Hoechst filter excluded compounds which were acutely toxic, auto-fluorescent, or which, through unknown mechanisms, affected Hoechst distribution. In summary, over four hundred 384-well plates were processed, and 123,450 compounds tested in the p-Ser65-Ub ICC phenotypic screen, with 15,197 identified hits, accounting for a primary hit rate of 12.3%. Stringent counter-screening and hit confirmation yielded a shortlist of 1,271 compounds and a final hit rate of 1.03% (Fig. 4F).

### Chemical and pharmacological profiles of hit compounds and determination of genotype- and stimulus-dependent compound activity

The 1,271 selected hit compounds were assessed for drug likeness (Fig. 5A-D). Molecular weight (Fig. 5A), number of hydrogen bond donors (Fig. 5B), LogP (Fig. 5C) and the high percentage passing Lipinski’s rule of 5 (Fig. 5D), all suggest that these compounds represent good starting points for further characterisation and hit-to-lead optimisation in a central nervous system (CNS)-targeting drug discovery program ^53–55^.

**Fig. 5.**
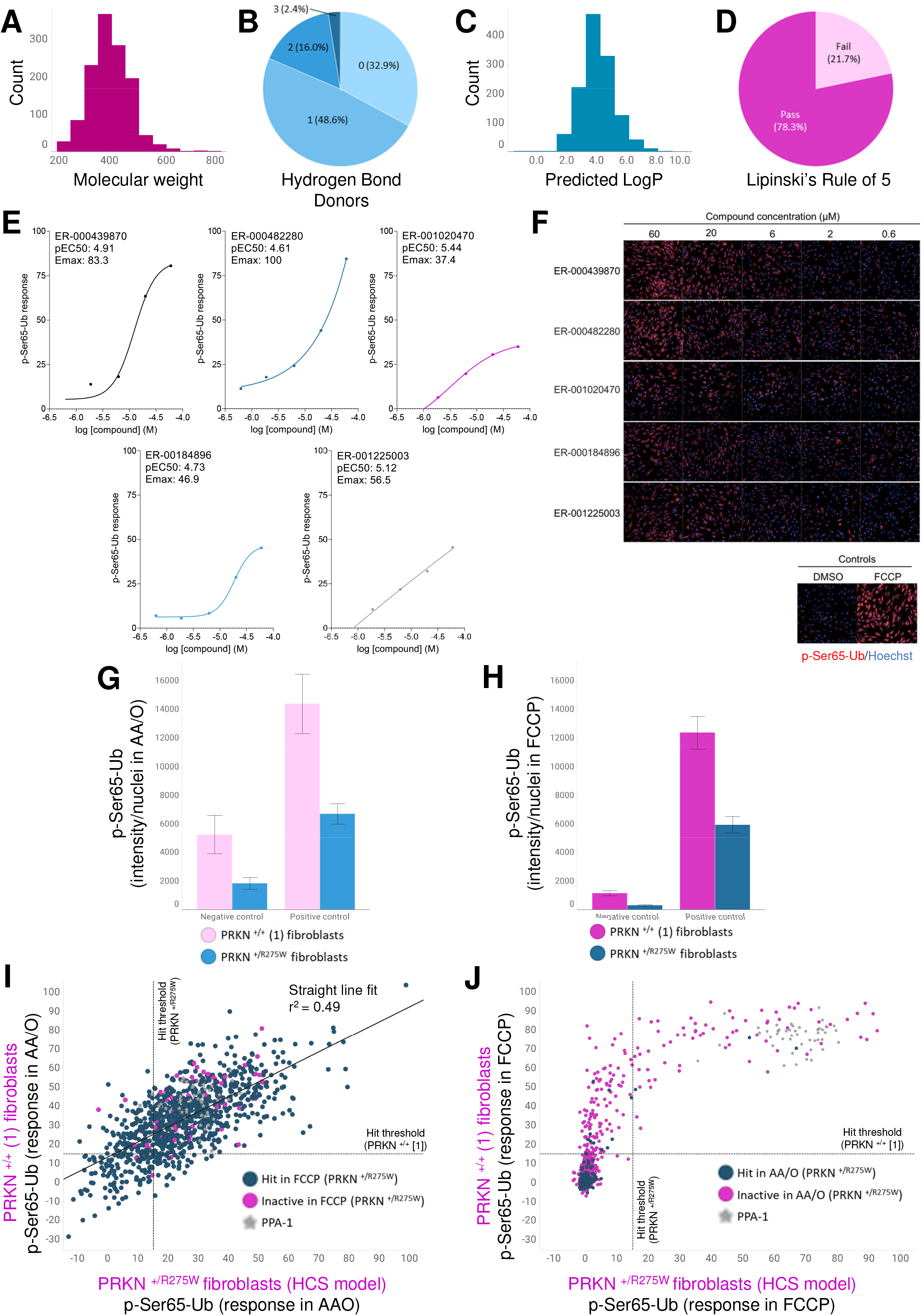
Chemical and pharmacological profiles of hit compounds and determination of genotype- and stimulus-dependent compound activity. **A**, Distribution of molecular weight of 1,271 hit compounds. **B**, Distribution of number of hydrogen bond donors of 1,271 hit compounds. **C**, Distribution of predicted partition coefficient (LogP) of 1,271 hit compounds, **D**, Distribution of 1,271 hit compounds which pass or fail Lipinski’s Rule of 5. **E**, Representative p-Ser65-Ub concentration-responses, Emax (maximal response within tested concertation range), and pEC50 (negative logarithm of the half maximal effective concentration), over five-point concentration curves (1:3 dilutions, 60 μM to 0.6 μM concentration range) using HCS conditions in Parkin ^+/R275W^ fibroblasts. **F,** Representative images (p-Ser65-Ub in red, Hoechst in blue) corresponding to compound concentration-response curves in **E**. **G**, p-Ser65-Ub response (p-Ser65-Ub intensity/nuclei, raw values) in Parkin mutant (Parkin ^+/R275W^) and control (Parkin +/+ (1)) human fibroblasts treated with DMSO (negative control) or FCCP (20 μM; positive control) under optimised HCS assay conditions (AA/O stimulus). **H,** p-Ser65-Ub response in Parkin mutant (Parkin ^+/R275W^) and control (Parkin +/+ (1)) human fibroblasts treated with DMSO (negative control) or FCCP (20 μM; positive control) using 2 h treatment with 2 μM FCCP as the stimulus (optimised HCS assay conditions for all other factors). **I,** Scatter plot and straight line fit of p-Ser65-Ub normalised response to test compounds using AA/O stimulus (HCS assay conditions), in Parkin mutant (Parkin ^+/R275W^; x-axis) and control (Parkin +/+ (1); y-axis) human fibroblasts. **J**, Scatter plot of p-Ser65-Ub normalised response to test compounds with 2 μM FCCP as the stimulus, in Parkin mutant (Parkin ^+/R275W^; x-axis) and control (Parkin +/+ (1); y-axis) human fibroblasts. For **G-H**, data are mean ± s.d. (n=96 (DMSO conditions) and n=48 (positive control) technical replicates). For **I-J**, grey stars indicate PPA-1 and number of tested compounds is 960.

To allow us to rank our hit compounds and further explore their effect on p-Ser65-Ub, we determined concentration-response relationships in the p-Ser65-Ub assay (Fig. 5E-F). Diverse concentration response profiles were present (examples in Fig. 5E-F). Maximal response ranged from 15.3 to 125.0 % normalised p-Ser65-Ub response, with the majority between 20 and 60%. Most compounds (about 75%) had a pEC50 between 4 and 5, representing an EC50 in the medium to high micromolar range, and about 10% of compounds had a pEC50 above 5, in the low micromolar range. Among the examples, ER-000439870 and ER-000482280 had high maximal responses, although ER-000482280 did not reach a plateau over the tested concentration range. ER-001020470, ER-000184896, and ER-001225003 had lower maximal responses, though ER-001020470 and ER-001225003 had high pEC50. We ranked compounds based on pEC50 for further investigation, to identify potential potent compounds of interest. While maximal response is also important, we have observed that high p-Ser65-Ub responses can correlate with MMP depolarisation (Fig. 4D), so we prioritised pEC50 to reduce the impact of depolarising compounds.

To further confirm compound activity, we tested selected hits in fibroblasts from a healthy control subject expressing common variant Parkin (PRKN ^+/+^; Fig. 5G-J), comparing an alternative mitophagy-inducing stimulus in place of AA/O, selecting the commonly used protonophore and mitochondrial membrane uncoupler FCCP at 2 μM (Fig. 5H and J). In the absence of test compound, the p-Ser65-Ub response to AA/O alone (0.5 μM AA) was approximately half in the Parkin ^+/R275^ compared to Parkin ^+/+^ fibroblasts (negative control; Fig. 5G). This genotype relationship was maintained following incubation with the positive control, 20 μM FCCP (Fig. 5G). Under mild FCCP (2 μM)-mediated mitochondrial stress alone, as well as in combination with the positive control (FCCP at 20 μM), the same genotype effect was also observed (Fig. 5H). Interestingly, p-Ser65-Ub intensity with 2 μM FCCP alone was reduced compared to 0.5 μM AA/O alone, suggesting that at these concentrations FCCP was less capable of inducing p-Ser65-Ub accumulation (Fig. 5G-H). However, addition of the positive control (FCCP at 20 μM) yielded a similar genotype dependent p-Ser65-Ub response, independent of initial stimulus (Fig. 5G-H), suggesting maximal pathway activation.

In the presence of AA/O-induced mitochondrial stress (HCS assay conditions), compound activity based on normalised p-Ser65-Ub response correlated well between Parkin ^+/R275W^ and Parkin ^+/+^ fibroblasts (Fig. 5I). The correlation indicates that the compound activity as a proportion of the positive control is relatively conserved and independent of genotype (Fig. 5I). In contrast, when using FCCP as the mitophagy stimulus instead of AA/O, hit compounds behaved differently between Parkin genotypes (Fig. 5J). Considerably fewer compounds hit overall, and the normalised p-Ser65-Ub response was dissimilar between the two cell lines. Instead, only compounds with >50% p-Ser65-Ub response in the control cells were also active in the Parkin ^+/R275W^ cells (Fig. 5J), suggesting a genotype dependency in these conditions. Despite the weak correlation across genotypes under mild FCCP observed for the test compounds, PPA-1 worked equally well between the two cell lines (grey stars in Fig. 5I-J).

### Functional and mechanistic insights into hit compound activity

p-Ser65-Ub accumulation reflects initiation of mitophagy and may not correlate with *en masse* mitochondrial degradation via the lysosome. To determine functional effects of our hit compounds on mitophagy, an eGFP clearance assay was established. Mitochondrially targeted-eGFP (mito-eGFP) was expressed transiently using a custom mRNA and expression was detected using high-content fluorescence imaging. Peak eGFP fluorescence occurred 20-24 hours post-transfection, following which a decay in eGFP signal intensity was observed. We demonstrated that mitophagy inducers, including AA/O, FCCP and valinomycin, increased the rate of decay of eGFP signal compared to DMSO (Fig. 6A; blue *versus* black), suggestive of clearance. Notably, co-incubation of bafilomycin A1 (BAF-A1), a vacuolar ATPase inhibitor which disrupts lysosomal function and autophagic flux, prevented the eGFP signal decay (Fig. 6A; pink), suggesting the decline in eGFP fluorescence after mitophagy induction is mediated via lysosomal/mitophagic degradation.

**Fig. 6.**
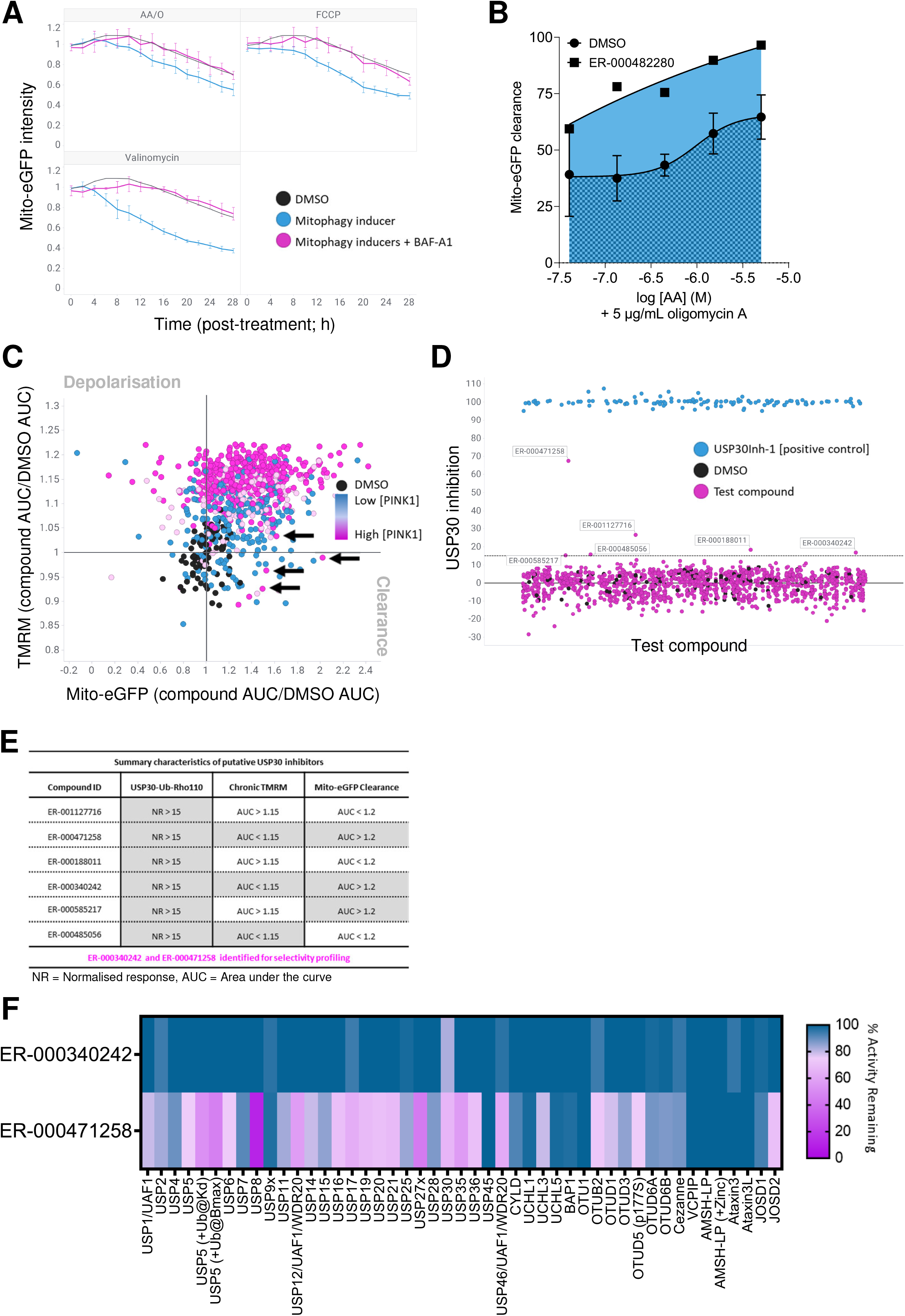
Functional and mechanistic insights into hit compound activity. **A**, Mito-eGFP intensity over a kinetic analysis of 28 hours, imaging at 2 h intervals, in the presence of mitophagy inducer (blue lines; 5 μM AA/O, 10 μM FCCP or 100 nM valinomycin. Bafilomycin A1 (BAF-A1, pink line) was included to inhibit lysosomal turnover of mito-eGFP (10 nM; 1 h pre-incubation). **B**, Representative compound displaying enhanced mito-eGFP clearance. A five-point concentration range of AA/O (0.04-5 μM) was used to induce mitophagy in cells subject to a single concentration of test compound or vehicle control (DMSO). Mito-eGFP fluorescence was recorded in live cells at 28 hours. Data normalised to DMSO (0% clearance) and valinomycin (200 nM; 100% clearance) without AA/O. **C**, Scatter plot of mito-eGFP clearance and TMRM response from 960 compounds, each run in singlicate. Hit compounds were pre-incubated for 2 h at 20 μM before addition of mitophagy inducing-stimulus and 1:2 dilution to a final assay concentration of 10 μM. A five-point concentration range of AA/O (0.04-5 μM) was used to induce mitophagy in cells subject to a single concentration of test compound or vehicle control (DMSO). Mito-eGFP fluorescence was recorded in live cells at 28 hours. Area under the five-point AA/O concentration curve (AUC) was calculated as normalised to assay controls valinomycin (200 μM) and DMSO without AA/O (100% and 0% mito-eGFP clearance respectively) and presented as fold change over the average DMSO AUC (derived from 6 technical DMSO replicates) on each assay plate. AUC was calculated based on the area between the fitted curve and the x-axis; test compound AUC is indicated by pale blue fill in **B**, DMSO AUC is indicated by dark blue check overlay, as indicated in **B**. Dots are coloured based on PINK1 AUC, derived as for mito-eGFP and TMRM. Gradient from low PINK1 (<1.2-fold change versus DMSO) in blue to high PINK (>1.4-fold change versus DMSO) in pink. **D,** Scatter plot of USP30-ubiquitin rhodamine 110 (Ub-Rho110) screen from ~1,200 compounds, to identify USP30 inhibitors. Compounds (including USP30Inh-1 positive control) pre-incubated at 20 μM for 30 minutes, before addition of 2xUb-Rho110 substrate. Normalised USP30 inhibition after 1 h reported (negative control, DMSO, 0% inhibition; positive control, USP30Inh-1, 100% inhibition). **E**, Summary characteristics of putative USP30 inhibitors over three assays. Grey shading indicates a positive outcome. TMRM and mito-eGFP data derived from **C**. **F**, DUB selectivity assay (DUBprofiler™) was conducted using 50 μM of ER-000340242 and ER-000471258. For **A**, data are mean ± s.d. (n=4 independent experiments). For **B**, data are mean ± s.d. (n=6 DMSO technical replicates). For **F**, data are mean of 2 technical replicates.

We then used our mito-eGFP clearance assay to assess hit compound-mediated mitochondrial clearance. We measured mito-eGFP clearance in live cells after 28 h compound treatment over a range of AA/O stimulus (exemplar compound, Fig. 6B; x-axis, Fig. 6C). A TMRM assay was carried out in the same samples following mito-eGFP acquisition to determine compound effects on MMP (Fig 6C; y-axis). To further multiplex this assay, following TMRM, the plate was processed for PINK1 ICC (Fig 6C; coloured circles). Leveraging the orthogonal read-outs, a range of compound profiles were identified from this single experiment. Interestingly, compounds with high MMP depolarisation also had high PINK1 signal, aligning with a known mechanism of PINK1 stabilisation and validating this approach. Notably, many compounds produced an increase in eGFP clearance (greater AUC) without perturbing MMP (Fig. 6C), with some even producing PINK1 stabilisation without MMP perturbation (Fig. 6C, black arrows).

Finally, as validation that our HCS and counter-screening approach yielded mechanistically relevant compounds, we searched within our 1,200 hits for inhibitors of the mitophagy negative regulator and deubiquitinase (DUB), USP30 (Fig. 6D). USP30 opposes mitochondrial substrate ubiquitination, modulating the threshold for mitophagy initiation ^56,57^. Inhibition of USP30 was assessed biochemically using the fluorogenic artificial DUB substrate, ubiquitin-rhodamine 110 (Ub-Rho110). Six compounds produced >15% inhibition of recombinant USP30 (Fig. 6D). Based on compound characterisation across both the USP30 activity and mito-eGFP functional assays (Fig. 6E), ER-000340242 and ER-000471258 were selected for profiling using the Ubiquigent DUBprofiler™ service. ER-000471258 demonstrated broad inhibitory effects against >40 known DUB enzymes at 50 μM, with very strong activity against USP8. In contrast, ER-000340242 proved more selective, with weak inhibitory effect (20-30% inhibition) only against USP30 (Fig. 6F).

## DISCUSSION

Defective mitochondrial quality control and accumulation of dysfunctional mitochondria are involved in the etiology of PD. Therefore, maintaining a healthy mitochondrial network through accelerating clearance of damaged mitochondria via mitophagy presents a therapeutic opportunity. Given the complexities of cellular signalling events, high-throughput screening (HTS)-compatible target-based biochemical assays, performed within an isolated or reconstituted system, may not represent the biology in a cellular context. Instead, in this study we have chosen a phenotypic screening approach, measuring p-Ser65-Ub accumulation. p-Ser65-Ub is a key upstream event in mitophagy initiation, reporting on activity of both PINK1 and Parkin, products of two PD-related genes, and is proven to be a key marker of mitochondrial quality control activation *in vivo*^58^. Together, the strong genetic rationale and clear cellular signalling events mediated by PINK1 and Parkin supported the development of an HCS-compatible assay.

We believe our phenotypic screening strategy has multiple advantages. (1) The HCS is mechanism-of-action and target-agnostic, likely to identify novel small molecule modulators of mitophagy; (2) the p-Ser65-Ub endpoint is pertinent to human PD genetics and relevant to the biology underlying disease pathophysiology (i.e., cellular ability to clear dysfunctional mitochondria); (3) due to its phenotypic nature it allows for identification of previously unknown molecular regulators of p-Ser65-Ub, not limited to canonical PINK1-Parkin-mediated mitophagy; and (4) it uses a cell system that is not based on Parkin overexpression, making it a more relevant physiological model. Together, the HCS may provide novel starting material for PD drug development, and a potent, selective small molecule therapeutic derived from this approach will likely have utility in sporadic PD.

To develop a robust high-content phenotypic screen measuring compound-induced changes in p-Ser65-Ub, we selected Parkin^+/R275W^ fibroblasts as a cellular model. These fibroblasts also have several advantages: (1) they manifest clear mitophagy deficits that can potentially be normalised; (2) despite these mitophagy deficits, the residual Parkin activity in these cells leaves them capable of accumulating p-Ser65-Ub (Fig. 1B) and allows the PINK1-Parkin pathway to be pharmacologically modulated (See ^38^ and Fig. 1B, Fig. 3); (3) fibroblasts are easily imaged and have clearly identifiable reticular mitochondria and (4) are readily expandable and amenable to HCS.

Selection of an appropriate cell model was followed by careful optimisation and validation of all aspects of the p-Ser65-Ub HCS assay (Fig. 1). We further developed a high-content TMRM assay for measurement of mitochondrial membrane potential (MMP; Fig. 2), to be used in counter-screening to remove compounds detrimentally affecting MMP. A proof-of-concept screen of forty-five selected literature compounds demonstrated that several compounds suggested to act via the PINK1-Parkin pathway were also active in our screening system, validating our approach (Fig. 3) ^43–46^.

We successfully screened a defined subset of the Eisai small molecule compound collection and identified molecules with the capacity to enhance p-Ser65-Ub accumulation in Parkin ^+/R275W^ patient-derived fibroblasts (Fig. 4). Following rigorous counter-screening, a final hit rate of 1% was determined, corresponding to our prediction in optimisation studies (Fig. 2). Hits were pharmacologically profiled across a five-point concentration range allowing further hit confirmation and prioritisation, and activity was confirmed in Parkin common variant fibroblasts (Fig. 5). We developed a mito-eGFP clearance assay to report on compound-induced functional clearance of mitochondrially-localised protein. Functional confirmation of hit compound activity was successfully multiplexed with chronic TMRM assessment and determination of compound-mediated PINK1 stabilisation: all three endpoints were determined over a concentration range of mitophagy-inducing stressor (Fig. 6). We have used these endpoints to prioritise and further shortlist our hit compounds.

As an example of a target-based deconvolution approach, we screened our hit compounds for USP30 inhibition, a proposed mechanism for mitophagy enhancement ^56^. We have previously observed that USP30Inh-1, 2 and 3 increase FCCP-mediated p-Ser65-Ub accumulation ^38^, and we further confirmed this in our proof-of-concept screen, observing a concentration-dependent increase in p-Ser65-Ub with these compounds, in the absence of mitochondrial membrane depolarisation (Fig. 3). Using an established USP30 biochemical assay ^38^, we identified several putative USP30 inhibitors, one of which (ER-000340242) proved selective across a panel of DUB enzymes (Fig. 6). These data provide confirmation that our HCS can yield mechanistically relevant biology and compounds. Further, comprehensive target-based and unbiased deconvolution approaches are required to understand compound mechanism-of-action of remaining hits.

One interesting observation was the dependence of hit compound activity as determined by the mitophagy-inducing stimuli across *PRKN* genotypes. We confirmed strong correlation between hit compound activity in *PRKN* common variant fibroblasts using the HCS mild AA/O stimulus, suggesting no genotype dependency (Fig. 5). However, using FCCP as an alternative mitophagy-inducing stimulus revealed a strong genotype effect. Hit compounds were more able to induce p-Ser65-Ub in *PRKN* common variant fibroblasts compared to Parkin ^+/R275W^ cells. One explanation for this may be that FCCP, at this low concentration, is less able to trigger mitophagy initiation and stabilise PINK1 in Parkin mutant cells. This is supported by the observation that the response to mild AA/O is greater in both cell lines compared to FCCP (Fig. 5G-H). It may be the threshold for mitophagy initiation has not been reached sufficiently in the *PRKN* mutant cells, given the feed-forward mechanism of Parkin function, and greater time or FCCP concentration is required. The dependence of hit compound activity on the mitophagy-inducing stress used (Fig. 5) emphasises the future value of disease-relevant *in vitro* models more closely reflecting pathophysiological stress.

In summary, we have successfully miniaturised, optimised and completed a phenotypic HCS using the p-Ser65-Ub endpoint in Parkin ^+/R275W^ fibroblasts. Through a well-defined workflow, we have identified compounds worthy of further investigation, which show activity in a novel functional assay, and which include modulators of the known mitophagy regulator USP30. Deconvolution and compound assessment in physiological and disease-based assays has the potential to identify novel biological regulators of PINK1-Parkin biology and provide starting material for drug discovery.

## MATERIALS AND METHODS

### Materials and compound synthesis

Compounds tested in the High Content Screen (HCS) were derived from the Eisai compound library. All other chemicals and compounds were purchased from Merck unless otherwise specified. All cell culture media and supplements were purchased from Thermo Fisher Scientific unless otherwise specified. Putative Parkin activators (PPA_s_) were derived from patent ID: WO 2018/023029 and synthesised in-house; structures are presented in Supplementary Table 2. Details of the compounds included in the proof-of-concept study (Fig. 3) can be found in Supplementary Table 3.

### Antibodies

For indirect immunocytochemistry (ICC), the following antibodies were used: mouse anti-PINK1 (Novus Biologicals, NBP236488, 1:1000). For direct ICC the following conjugated antibodies were used: Alexa Fluor 647-labelled anti-p-Ser65-Ub (Cell Signalling Technology, 62802, 1:1500), Alexa Fluor 488-labelled anti-HSP60 (Abcam, ab128567, 1:1000), and Alexa Fluor 555 labelled-anti-LAMP1 (BD Biosciences, 555798, 1:1000). Anti-p-Ser65-Ub was conjugated by CST using their basic conjugation service. Anti-HSP60 and anti-LAMP1 were conjugated using Lightning Link Alexa 488 (Abcam, ab236553) and Alexa 555 (Abcam, ab269820) respectively, following the manufacturer’s protocol, with an overnight incubation at room temperature.

### Cell culture

Control and patient-derived fibroblasts were obtained from the NINDS Human Cell and Data Repository (https://stemcells.nindsgenetics.org). Two cell lines were primarily used: Parkin^+/+^ unaffected healthy control (ND36320; labelled as ‘+/+ (1)’ in the fibroblast characterisation panel in Fig. 1B) and Parkin ^+/R275W^ (ND29369; ‘+/R275W’; used for the HCS). The following additional fibroblast lines were used (Fig. 1B): Parkin ^+/+^ (ND34769 [+/+ (2)]), Parkin ^+/+^ (ND34770 [+/+ (3)]), Parkin ^N52fs/Ex3-4Del^ (ND29543), Parkin ^R42P/Ex3Del^ (ND30171), Parkin ^R42P/+^ (ND31618), Parkin ^Ex4-7Del/Q43fs^ (ND40067), Parkin^R275W/R275Q^ (ND40078), and Parkin ^R245K/G430D^ (NN0004771/NH50289). These fibroblast lines are summarised in Supplementary Table 1. All fibroblasts were sequenced for *PRKN* (Genewiz), verifying the above genotypes. Cell expansion to provide sufficient biomass for the HCS was performed by SAL Scientific.

Fibroblasts were grown in Dulbecco’s Modified Eagle Medium, high glucose, with GlutaMAX (DMEM; Gibco, 61965-026), containing 10% fetal bovine serum (FBS; Biosera, FB1350-500), 1 mM sodium pyruvate (Gibco, 11360) and 100 U/mL penicillin/100 μg/mL streptomycin (Gibco, 15140). For galactose metabolism experiments, fibroblasts were grown in Dulbecco’s Modified Eagle Medium with no glucose, no glutamine, no phenol red (Gibco, 12307263), containing 4.5g/L galactose (Merck, G-0625), GlutaMAX (Fisher, 11574466), 10% fetal bovine serum (dialysed, US origin, One Shot™, Fisher, A3382001), 1 mM sodium pyruvate and 100 U/mL penicillin/100 μg/mL streptomycin.

Human neuroblastoma control SH-SY5Y cell line (Abcam, ab275475), Parkin knockout (*PRKN*^−/−^) SH-SY5Y cells (Abcam, ab280101) and PINK1 knockout (*PINK1*^−/−^) SH-SY5Y cells (Abcam, ab280876) were cultured in Dulbecco’s Modified Eagle Medium/F-12 (1:1) with GlutaMAX (DMEM/F-12; Gibco 31331-028), supplemented with 10% FBS (Biosera, FB1350-500) and 100 U/mL penicillin/100 μg/mL streptomycin.

### HCS assay: p-Ser65-Ub direct ICC

p-Ser65-Ub was assessed using direct ICC. Freshly thawed patient-derived fibroblasts were seeded at 3000 cells per well in 25 μL pre-warmed complete cell culture medium in CellCarrier Ultra 384 well plates (PerkinElmer, 6057300) using a Multidrop Combi Reagent Dispenser (Thermo Scientific), left to adhere at room temperature for 1 h, then incubated overnight at 37°C and 5% CO_2_. Medium was replaced with 25 μL of pre-warmed complete cell culture medium containing compounds using the CyBio Cybi Well Vario (Analytik Jena) and incubated for 2 h at 37°C and 5% CO_2_. A further 25 μL of pre-warmed complete cell culture medium containing mitochondrial toxins at twice the final assay concentration (FAC; FAC of 0.5 μM antimycin A, 5 μg/mL oligomycin A) was added using the CyBio Cybi Well Vario and incubated for a further 2 h. Any deviations from these conditions are defined within figure legends. In AA/O treatments, oligomycin A was always used at a FAC of 5 μg/mL.

Medium was removed and cells were fixed in ice-cold acetone: methanol (A:M; 1:1) for 30 seconds, added using the CyBio Cybi Well Vario. A:M was removed and 25 μL of ice-cold D-PBS (without Ca^2+^/Mg^2+^) was added using the Multidrop Combi Reagent Dispenser, then incubated on ice for 30 minutes. D-PBS was replaced with ice-cold blocking solution (3% BSA [Sigma, A3803] in D-PBS, without Ca^2+^/Mg^2+^) using a Dragonfly Discovery liquid dispenser (SPT Labtech) and incubated for 1 h on ice. Blocking solution was replaced with ice-cold blocking solution (3% BSA in D-PBS) containing conjugated antibodies (Alexa 647 anti-p-Ser65-Ub [CST, 62802, 1:1500], Alexa 488 anti-HSP60 [Abcam, ab128567, 1:1000], and Alexa 555 anti-LAMP1 [BD Biosciences, 555798, 1:1000]) using a Dragonfly Discovery liquid dispenser and incubated at 4°C overnight in the dark with gentle agitation.

Cells were washed twice in room temperature D-PBS containing 0.1% Tween 20 (D-PBS-T) using the Multidrop Combi Reagent Dispenser, then 25 μL of Fluoromount G (SouthernBiotech, 0100-01): D-PBS (1:1) with Hoechst (2 μg/mL) was added per well using the Dragonfly Discovery liquid dispenser. Plates were sealed using opaque foil seals (Agilent, 24214-001) and incubated for at least 15 minutes at room temperature. Images were acquired using the Opera Phenix (PerkinElmer) using the 20x water objective (NA 1.0, WD 1.7 mm, field of view approximately 646 μm x 646 μm) and 4 fields imaged per well.

### p-Ser65-Ub direct ICC in SH-SY5Y cells

p-Ser65-Ub was assessed in SH-SY5Y cells using direct ICC as detailed for the HCS assay but with minor modifications. SH-SY5Y cells were seeded at 12,500 cells per well in pre-warmed complete cell culture medium in CellCarrier Ultra 384 well plates using the Dragonfly Discovery liquid dispenser and incubated overnight at 37°C and 5% CO_2_. Medium was replaced with 25 μL of pre-warmed complete cell culture medium containing treatments using the CyBio Cybi Well Vario and incubated for 2 h at 37°C and 5% CO_2_. In AA/O treatments, oligomycin A was always used at a final assay concentration of 5 μg/mL.

Medium was removed and cells were fixed in ice-cold A:M (1:1) for 30 seconds, added using the CyBio Cybi Well Vario. A:M was removed and 25 μL of ice-cold D-PBS (without Ca^2+^/Mg^2+^) was added using the Cybio Cybi Well Vario, then incubated on ice for 30 minutes. D-PBS was replaced with ice-cold blocking solution (3% BSA in D-PBS) using the Dragonfly Discovery liquid dispenser and incubated for 1 h on ice. Blocking solution was replaced with ice-cold blocking solution (3% BSA in D-PBS) containing all conjugated antibodies (Alexa 647 anti-p-Ser65-Ub [CST, 62802, 1:1500], Alexa 488 anti-HSP60 [Abcam, ab128567, 1:1000], and Alexa 555 anti-LAMP1 [BD Biosciences, 555798, 1:1000]) using the Dragonfly Discovery liquid dispenser and incubated at 4°C overnight in the dark with gentle agitation.

Cells were washed twice in room temperature D-PBS with 0.1% Tween 20 (D-PBS-T) the Cybio Cybi Well Vario, then 25 μL of Fluoromount G: D-PBS (1:1) with Hoechst (2 μg/ml) was added per well using the Dragonfly Discovery liquid dispenser. Plates were sealed using opaque foil seals and incubated for at least 15 minutes at room temperature before image acquisition. Images were acquired using the Opera Phenix using the 20x water objective (NA 1.0, WD 1.7 mm, field of view approximately 646 μm x 646 μm) and 4 fields imaged per well.

### Mitochondrial membrane potential assay: TMRM

Mitochondrial membrane potential was measured using tetramethyl rhodamine methyl ester (TMRM) in re-distribution mode ^38^. Freshly thawed patient-derived fibroblasts were seeded at 3000 cells per well in CellCarrier Ultra 384 well plates, left to adhere at room temperature for 1 h, then incubated overnight at 37°C and 5% CO_2_. Medium was replaced with 25 μL of pre-warmed complete cell culture medium containing compounds using CyBio Cybi Well Vario and incubated for 2 h. A further 25 μL of pre-warmed complete cell culture medium containing mitochondrial toxins (FAC 0.5 μM antimycin A, 5 μg/mL oligomycin A) in TMRM staining solution (FAC 50 nM TMRM, 1 μg/mL Hoechst) was added using CyBio Cybi Well Vario and incubated for 2 h at 37°C to allow equilibration of dye. In AA/O treatments, oligomycin A was always used at a final assay concentration of 5 μg/mL. Images were acquired using the Opera Phenix, 20x water objective (NA 1.0, WD 1.7 mm, field of view approximately 646 μm x 646 μm), 2 fields imaged per well, with temperature and CO_2_ controls enabled.

### Functional mitophagy assay: mito-eGFP clearance

Fibroblasts were transfected using custom mRNA, synthesised *de novo* (provided by Trilink Biotechnologies). Mitochondrial targeted-enhanced Green Fluorescent Protein (Mito-eGFP) mRNA was designed by appending a N-terminal mitochondrial localisation sequence derived from COX8a (accession number: NP_004065.1) to eGFP. 25 μg mRNA was transfected into 2.5 × 10^6^ freshly thawed fibroblasts using Cell Line Nucleofector Kit V (Lonza) and AMAXA program X-001 following the manufacturer’s instructions. Cells were seeded at 5000 cells per well in CellCarrier Ultra 384 well plates, left to adhere at room temperature for 1 h, then incubated overnight at 37°C and 5% CO_2_.

Compounds were added in a full medium change at 20 h following transfection and cell seeding. Medium was replaced with 25 μL of pre-warmed complete cell culture medium containing compounds using the CyBio Cybi Well Vario and incubated for a further 2 h. A further 25 μL of pre-warmed complete cell culture medium containing mitochondrial toxins at twice the FAC (5-point concentration range of antimycin A [FAC 0.04-5 μM] with oligomycin A [constant FAC 5 μg/mL] and Hoechst [FAC 1 μg/mL]) was added using CyBio Cybi Well Vario and incubated for 28 h. eGFP signal was acquired using the Opera Phenix, 20x water objective (NA 1.0, WD 1.7 mm, field of view approximately 646 μm x 646 μm), 3 fields imaged per well, with temperature and CO_2_ controls enabled.

Following imaging of eGFP, a further 5 μL of pre-warmed complete cell culture medium containing 6x TMRM staining solution (300 nM TMRM; FAC 50 nM) was added using the Dragonfly Discovery liquid dispenser and incubated for 1 h at 37°C to allow equilibration of dye. Images were acquired using the Opera Phenix, 20x water objective (NA 1.0, WD 1.7 mm, field of view approximately 646 μm x 646 μm), 2 fields per well, with temperature and CO_2_ controls enabled. Following live-cell imaging, cells were fixed and immuno-stained for PINK1, following the indirect ICC method below.

### Indirect ICC

For fibroblasts, medium was removed, and cells were fixed in ice-cold A:M (1:1) for 30 seconds, added using CyBio Cybi Well Vario. A:M was removed and 25 μL of ice-cold D-PBS (without Ca^2+^/Mg^2+^) was added using Multidrop Combi Reagent Dispenser, then incubated on ice for 30 minutes. D-PBS was replaced with ice-cold blocking solution (3% BSA in D-PBS) using the Dragonfly Discovery liquid dispenser and incubated for 1 h on ice. Blocking solution was replaced with ice-cold blocking solution (3% BSA in D-PBS) containing appropriate antibodies using the Dragonfly Discovery liquid dispenser and incubated at 4°C overnight with gentle agitation.

Cells were washed three times in room temperature TBS with 0.1% Tween 20 (TBS-T) using Multidrop Combi Reagent Dispenser, then room temperature blocking solution (3% BSA in D-PBS) containing secondary antibody (IgG2b Cross-Adsorbed Goat anti-Mouse, Alexa Fluor™ 555 [Invitrogen, 10412832], 1:2000) was added using Dragonfly Discovery liquid dispenser and incubated for 1 h at room temperature. Cells were again washed three times in room temperature TBS-T using the Multidrop Combi Reagent Dispenser, then 25 μL of Fluoromount G: TBS (1:1) with Hoechst (2 μg/mL) was added per well using the Dragonfly Discovery liquid dispenser. Plates were sealed using opaque foil seals and incubated for at least 15 minutes at room temperature before image acquisition. Images were acquired using the Opera Phenix, 40x water objective (NA 1.1, WD 0.62 mm, field of view approximately 323 μm x 323 μm), 8 fields imaged per well.

### USP30-Ub Rho110 activity assay

USP30-UbRho10 was performed as previously described ^38^. Compounds were dispensed into black, clear-bottom, 384 well plates (Greiner, 781096) using the ECHO 550 (Labcyte) liquid handler. 2× FAC His-tagged recombinant human USP30 protein (rhUSP30; amino acids 57–517 of the full-length protein, and a C-terminal 6-His tag, Sf 21 (baculovirus)-derived; 10 nM, FAC 5 nM [R&D Systems, E-582-050]) was prepared in USP30 activity assay buffer (50 mM Tris base pH 7.5, 100 mM NaCl, 0.1 mg/ml BSA [Sigma, A7030], 0.05% Tween 20, 1 mM DTT) and 15 μL dispensed into compound-containing assay plate using the Dragonfly Discovery liquid dispenser and incubated for 30 minutes at room temperature. Following incubation, 15 μL of 2x concentrated ubiquitin-rhodamine 110 (Ub-Rho110; 200 nM, FAC 100 nM [R&D Systems, U-555-050]) in USP30 activity assay buffer was dispensed into the compound-rhUSP30 containing plate using the Dragonfly Discovery liquid dispenser and fluorescence read on the FLIPR TETRA plate reader (Molecular Devices). Fluorescence intensity was recorded over 1 h and normalised to positive and negative controls (USP30Inh-1 ^38^, 10 μM and DMSO respectively).

### Deubiquitinase selectivity profiling

Selectivity profiling against forty-one deubiquitinase (DUB) enzymes was performed at Ubiquigent (Dundee, U.K.) using the DUBprofiler™ platform and Ub-Rho110-glycine substrate based-assay.

### Data analysis and statistics

Data are presented as mean ± standard deviation (SD). Normalisation of the data allowed for control of inter-assay variability. Assay quality was determined after calculation of Z’, using the equation: 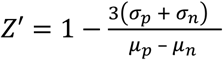, with standard deviation (σ) and means (μ) of the positive (p) and negative (n) controls ^39^.

For the HCS p-Ser65-Ub ICC assay, image analysis was performed using the Perkin Elmer Harmony PhenoLOGIC machine learning plug-in, which was used to score cells as either p-Ser65-Ub positive or p-Ser65-negative, based on the image texture phenotype of positive and negative controls. PhenoLOGIC plug-in enables the recognition of phenotypes within cell populations using a machine learning approach that combines the most meaningful parameters to achieve accurate classification of cell phenotypes at a single cell level. Data is presented as percentage positive cells normalised to positive and negative controls (FCCP, 20 μM FAC, and DMSO respectively). Plates with Z’ ≥ 0.8 and secondary QC ≥ 20% were considered to pass QC. For the TMRM assay, Z’≥ 0.8 was considered a pass. For all other high-content imaging, including hit compound pharmacology in p-Ser65-Ub assay, fluorescence intensity of signal was calculated per nuclei number and normalised to positive and negative controls.

Data handling, statistical analysis, and data visualisation were performed using Graphpad Prism 9 and Tibco Spotfire Analyst 10.3.3. Curve fitting was performed using the IDBS XLfit (5.4.0.8) add-in for Microsoft Excel. Statistical tests are indicated in figure legends. Statistical significance was assessed as being P < 0.05.

For the mito-eGFP functional mitophagy assay, data were normalised to positive and negative controls (200 μM valinomycin and DMSO respectively). Area under the curve (AUC) over the 5-point AA/O concentration range was calculated for both test compound and 6x DMSO technical replicates per plate using the IDBS XLfit (5.4.0.8) add-in for Microsoft Excel and presented as fold change relative to average DMSO AUC (compound AUC/DMSO AUC).

Chemical profile of the selected hit compounds was generated by ChemAxon v16.10.24

## ACKNOWLEDGEMENTS

We thank Robin Ketteler and Eliona Tsefou (University College London) for their input into the project. We are grateful to the Michael J Fox Foundation (MJFF Grant ID: MJFF-009659) for generously funding a portion of this study. We would like to thank members of the UCL: Eisai Therapeutic Innovation Group (Andy Takle, Tom Warner, Peter Atkinson, Hélène Plun-Favreau, Adrian Isaacs, Anthony Groom, Jane Kinghorn and Nicola Ridgway) for scientific discussions, guidance, and critique of the project. Finally, we thank Aurelio Vázquez de la Torre for his contribution in the early stages of this project.

## AUTHOR CONTRIBUTIONS

C.T. and J.S. performed and analysed experiments on the *PRKN* mutant fibroblast panel. R.T., E.C., and T.B. performed and analysed all other experiments, including the HCS assay optimisation and primary screening. T.H. and K.T. provided chemistry support throughout the project, including identification and synthesis of the putative Parkin activators. R.T., E.C., and T.B. wrote the manuscript. J.M.S., T.H. and C.T. edited the manuscript. T.B. and J.M.S. conceived the project. T.B. supervised the project.

