## Supplementary Tables for "High-content phenotypic screen to identify small molecule enhancers of Parkin-dependent ubiquitination and mitophagy"

**Supplementary Table 1.** Human primary skin fibroblast panel

| **Parkin status** | **Cell line ID (NINDS)** | **Clinical status** | **Age** | **Gender** |
|---|---|---|---|---|
| +/+ (1) | ND36320 | unaffected | 71 | F |
| +/+ (2) | ND34769 | unaffected | 68 | F |
| +/+ (3) | ND34770 | unaffected | 72 | M |
| +/R275W | ND29369 | parkinsonism | 61 | F |
| R275W/R275Q | ND40078 | parkinsonism | 51 | F |
| R42P/Ex3 DEL | ND30171 | parkinsonism | 54 | M |
| +/R42P | ND31618 | parkinsonism | 63 | F |
| N52fs//Ex3-4 DEL | ND29543 | parkinsonism | 50 | M |
| Ex4-7 DEL/Q43fs | ND40067 | parkinsonism | 44 | F |
| R245K/G430D | NN0004771/ NH50289 | dystonia | 38 | M |

**Supplementary Table 2.** Putative Parkin activator structures

| | |
|---|---|
| **PPA-1** | 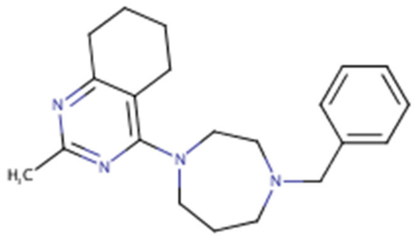 |
| **PPA-2** | 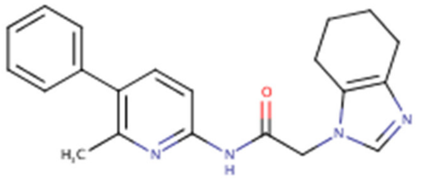 |
| **PPA-3** | 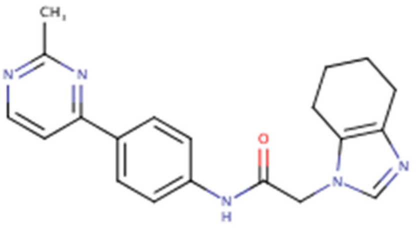 |
| **PPA-4** | 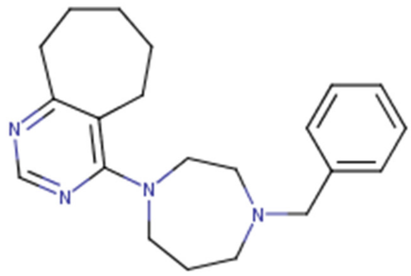 |

**Supplementary Table 3.** Reference compounds

| Reference molecule | Starting (stock) concentration (mM) | Supplier | Cat number |
|---|---|---|---|
| A-769662 | 10 | Merck | SML2578 |
| Antimycin A (AA) | 5 | Merck | A8674 |
| Antimycin A / Oligomycin A (AA/O) | 5/10 mg/mL | Merck/Sellek | A8674/S1478 |
| Actinonin | 100 | Merck | A6671 |
| AICAR | 10 | Merck | A9978 |
| Ambroxol | 20 | Merck | A9797 |
| Bafilomycin A1 | 0.01 | Merck | B1793 |
| BAM15 | 10 | Merck | SML1760 |
| BAY 87-2243 | 10 | Merck | SML2384 |
| BC1464 | 10 | In-house | NA |
| Bz-423 | 10 | Merck | SML1944 |
| Conduritol B epoxide (CBE) | 10 | Merck | C5424 |
| CCCP | 10 | Merck | C2759 |
| Compound 3 3-(3-Fluorophenyl)-1-(4-{[6-(1H-imidazol-1-yl)pyrimidin-4-yl]amino}phenyl)urea) | 10 | Chembridge | 9211821 |
| Cyclosporin A | 10 | Merck | C3662 |
| Deferiprone | 10 | Merck | Y0001976 |
| DMSO | NA | Merck | D8418 |
| ER-000444793 | 10 | In-house | NA |
| FCCP | 10 | Merck | C2920 |
| Kinetin | 10 | Merck | K3378 |
| Metformin | 10 | Merck | PHR1084 |
| MPP+ | 500 | Merck | D048 |
| MWP00839 | 10 | Aurora Fine Chemicals LLC | K02.225.465 |
| Nicotinamide | 100 | Merck | N0636 |
| Nilotinib | 10 | Merck | CDS023093 |
| Oligomycin | 10 mg/mL | Sellek | S1478 |
| Pifithrin a | 10 | Merck | P4359 |
| PMI | 10 | Probechem | PC-62488 |
| PPA-1 | 10 | In-house | NA |
| PPA-2 | 10 | In-house | NA |
| PPA-3 | 10 | In-house | NA |
| PPA-4 | 10 | In-house | NA |
| Rapamycin | 10 | Millipore | 553210 |
| Rotenone | 10 | Merck | R8875 |
| SMER28 | 10 | Merck | S8197 |
| SPB08007 | 10 | Aurora Fine Chemicals LLC | K13.701.129 |
| SR3677 | 10 | Merck | SML0774 |
| T0466/GW806742X | 10 | Merck | SML1990 |
| T-271 | 10 | In-house | NA |
| Thapsigargin | 10 | Merck | T9033 |
| TORIN1 | 1 | Tocris | 4247/10 |
| Urolithin A | 10 | Merck | SML1791 |
| USP30Inh-1 | 10 | In-house | NA |
| USP30Inh-2 | 10 | In-house | NA |
| USP30Inh-3 | 10 | In-house | NA |
